# *Wolbachia* confers sex-specific resistance and tolerance to enteric but not systemic bacterial infection in *Drosophila*

**DOI:** 10.1101/045757

**Authors:** Radhakrishnan B. Vasanthakrishnan, Gupta Vanika, Jonathon Siva-Jothy, Katy M. Monteith, Sam P. Brown, Pedro F. Vale

## Abstract

*Wolbachia*-mediatedprotection against viral infection has been extensively demonstrated in *Drosophila* and in mosquitoes that are artificially inoculated*with D. melanogaster Wolbachia* (wMel), but to date no evidence for *Wolbachia*-mediated antibacterial protection has been demonstrated in *Drosophila.*Here we show that *D. melanogaster* carrying wMel shows reduced mortality during enteric – but not systemic - infection with the opportunist pathogen *Pseudomonas aeruginosa*, and that protection is more pronounced in male flies. *Wolbachia-mediated* protection is associated with increased early expression of the antimicrobial peptide *attacinA*, followed by increased expression of a ROS detoxification gene (*gstD8*), and other tissue damage repair genes which together contribute to greater host resistance and disease tolerance. These results highlight that the route of infection is important for symbiont-mediated protection from infection, that *Wolbachia* can protect hosts by eliciting a combination of resistance and disease tolerance mechanisms, and that these effects are sexually dimorphic.

## Introduction

All organisms experience a combination of beneficial and detrimental colonisations by pathogens, commensals and symbionts, with profound effects on host physiology, behaviour, ecology and evolution (Bennett and Moran, 2015; Douglas, 2015; Gandon and Vale, 2014; Lewis and Lizé, 2015; Werren et al., 2008). Bacterial endosymbionts of insects, for example, are known to manipulate host reproduction (Engelstädter and Hurst, 2009; Werren et al., 2008), to alter the host's acquisition of essential nutrients(Douglas, 2015, 1998), and to provide protection from the deleterious effects of parasites and pathogens (Brownlie and Johnson, 2009; Hamilton and Perlman,2013).

*Wolbachia* is a maternally-inherited, intracellular bacterium of arthropods and nematodes, and is one of the best studied microbial symbionts (Brownlie and Johnson, 2009; Hamilton and Perlman, 2013). Its host range is vast, with recent estimates that 48-57% of all terrestrial arthropods (Weinert et al., 2015), and at least 10% of all *Drosophila* species carry *Wolbachia* (Mateos et al., 2006). The ability of some *Wolbachia* strains to protect insect hosts from pathogenic infections make it particularly relevant for potential bio-control of insect vectored zoonotic infections, and more broadly, relevant as mediators of pathogen-mediated selection in insects (Brownlie and Johnson, 2009; Hamilton and Perlman, 2013; Karyn N. Johnson, 2015). *Aedes aegypti* and *Ae. albopictus* mosquitoes, for example, have been shown to become more resistant to Dengue and Chikungunya viruses, as well as malaria-causing *Plasmodium* when they are experimentally inoculated with *Wolbachia* (Bian et al., 2010; Kambris et al., 2010; Moreira et al., 2009). In *Drosophila*, there is also ample evidence that flies carrying *Wolbachia* are better able to survive infection by a number of naturally occurring RNA viruses (Hedges et al., 2008; Hedges and Johnson, 2008; Karyn N Johnson, 2015; Teixeira et al., 2008). This anti-viral protection is variable among strains of *Wolbachia* and correlates strongly with the reduction in viral titres within hosts (Martinez et al.,2014), suggesting that *Wolbachia* generally enhances the ability to clear pathogens (increasing host resistance) rather than the ability to repair damage independently of pathogen clearance (disease tolerance) (Ayres and Schneider, 2012; Råberg et al., 2009).

In contrast to the strong evidence for *Wolbachia*-mediated antiviral protection, its ability to protect its native fruit fly hosts from bacterial infections has not been clearly demonstrated to date. In one study, carrying *Wolbachia* did not affect the survival or immune activity of *D. simulans* or *D. melanogaster* during systemic infection with *Pseudomonas aeruginosa, Serratia marcescens* or *Erwinia carotovora* (Wong et al., 2011), while another study found that the presence of *Wolbachia* had no effect on the ability to suppress pathogen growth during systemic infections by intracellular (*Listeria monocytogenes, Salmonella typhimurium*) or extracellular bacterial pathogens (*Providencia rettgeri*) (Rottschaefer and Lazzaro, 2012). Given that *Wolbachia* can provide broad-spectrum protection against a range of pathogens, including bacteria, in mosquitoes (Ye et al., 2013), the lack of evidence for antibacterial protection in flies is puzzling. Some authors have proposed that antibacterial protection may only occur in novel host-*Wolbachia* associations (like those of mosquitoes), although the exact mechanism for such protection remains unclear (Wong et al., 2011; Zug and Hammerstein, 2015).

One possibility is that the experimental conditions under which *Drosophila* are commonly challenged with pathogens in the lab do not reflect the infections they are likely to encounter in the wild. For example, experimental infections often focus on systemic infection, introducing large quantities of bacterial pathogens via intra-thoracic or abdominal injection (Neyen et al., 2014; Rottschaefer and Lazzaro, 2012; Wong et al.,2011). The ecological context of fruit flies however, which consists mainly of foraging on rotting organic matter, means that most pathogens in the wild are more likely to be acquired orally, resulting in enteric, rather than systemic infections (Ferreira et al., 2014; Stevanovic and Johnson, 2015). It is therefore conceivable that any form of *Wolbachia*-mediated protection that could have evolved in the context of enteric infection may not be detected during systemic infection.

Here we show that the route of infection is indeed important for *Wolbachia*-mediated protection in *Drosophila*, which we find to occur during enteric - but not systemic - infection by the opportunist pathogen *Pseudomonas aeruginosa. P. aeruginosa* has an incredibly broad host range, infecting insects, nematodes, plants, and vertebrates, and is found in most environments (Apidianakis and Rahme, 2009; Neyen et al., 2014). Enteric infection of *Drosophila* by *P. aeruginosa* results in pathology to intestinal epithelia due the formation of a bacterial biofilm in the crop, a food storage organ in the foregut (Mulcahy et al., 2011; Sibley et al., 2008). In the majority of enteric infections *P. aeruginosa* growth is restricted to the crop, and is sufficient to cause death (Chugani et al., 2001; Sibley et al., 2008). We exposed flies that were naturally infected with *Wolbachia*, and identical derived flies that were cured of *Wolbachia* infection, to *P. aeruginosa* either through intra-thoracic pricking (causing a systemic infection) or through the oral route of infection by feeding (causing an enteric infection). We then monitored how within-host microbe loads and survival varied throughout the course of an infection to assess if (1) *Wolbachia*-mediated protection occurred during systemic and enteric bacterial infection; (2) when protection was detected, if this was due to differences in the bacterial clearance rate (resistance) or if it aided host survival despite high microbe loads (tolerance); and (3) how these protective effects differed between male and female flies. We further characterized the expression of immune and damage repair genes previously shown to be involved in enteric bacterial infection in *Drosophila*.

## Results

### *Wolbachia* reduces mortality during enteric but not systemic bacterial infection

All flies infected systemically with PA14 via intra-thoracic pricking died within 24 hours (Fig 1a), and in line with previous work (Wong et al., 2011), we did not detect any significant effect of *Wolbachia* status on the rate at which they died (Cox Proportional Hazard Model, Likelihood Ratio X^2^ = 0.003, DF=1, p=0.959), regardless of sex ( ‘Sex’ effect, X^2^= 0.860, DF=1, p=0.354). Flies that ingested and acquired an enteric infection of PA14 died at a faster rate than control flies exposed only to a sucrose solution (**Fig. 1b;** ‘Infection status’ effect, Likelihood Ratio X^2^= 64.27, DF=1, p<0.0001). Fly mortality during enteric infection was significantly affected by their *Wolbachia* status (X^2^= 6.32, DF=1, p=0.013). This protective effect was not substantial in female flies: female flies without *Wolbachia* were 1.58 more likely to die than infected females carrying *Wolbachia* (Cox risk ratio, X^2^ = 1.27, p=0.2644). The protection in male flies was more pronounced, where not carrying *Wolbachia* made PA14-infected males 2.26 times more likely to die than their infected *Wolbachia-positive* counterparts (Cox risk ratio, X^2^ = 4.22, p=0.0172)(Figure 1b). In order to understand the cause of the observed protection during enteric but not systemic infection protection, the results below focus only on flies that acquired infection orally.

**Figure 1.**
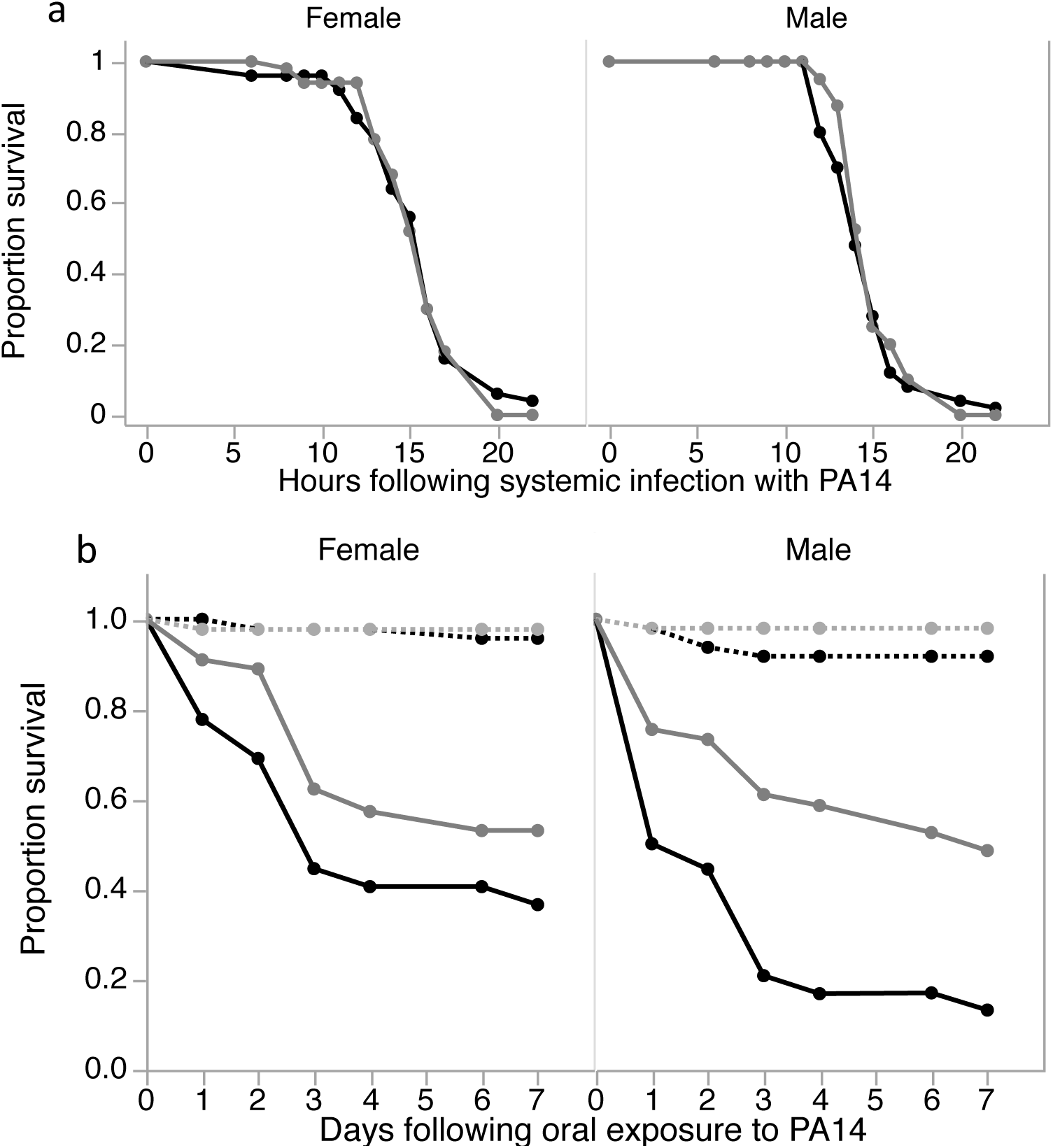
Fly survival after systemic (Fig. 1a) oral infection (Fig. 1b) with *P. aeruginosa*PA14. OreR^Wol-^ (black line), OreR^Wol+^ (grey line) were either pricked with a needle dipped in PA14 culture (OD = 1), or left to feed on a PA14 culture (OD=23) or on a control solution of 5% sugar for 12 hours. Survival was monitored for 24h (systemic infection) or daily (oral infection). Data were analysed using a Cox Proportional Hazard model.

### *Wolbachia* does not affect the rate of bacterial clearance during enteric infection

Following 12 hours of exposure to *P. aeruginosa*, bacterial loads decreased over the course of the experiment in both male and female flies (**Fig. 2**) time effect F7,186=48.81, p<0.0001). However, the rate at which infection was cleared was not affected by *Wolbachia* status (‘Time × Wolbachia’ interaction F7,186=5.71, p=0.30), which suggests that the presence of *Wolbachia* does not contribute to the clearance of this bacterial gut infection. Regardless of *Wolbachia* status, we observed that males and females showed different patterns of bacterial clearance over time (**Fig. 2;** ‘Time × Sex’ interaction F7,186=4.21, p=0.002). While males appeared to be able to clear the infection almost entirely within a week (mean ± SEM 0.85 ±0.29 Log10 CFU per fly at 168 hours post exposure) females appeared to stop clearing infection after 96h, maintaining a relatively stable bacterial load of about 100 CFUs per fly until the end of the experiment (**Fig. 2**). These sex differences in bacterial clearance were present regardless of the *Wolbachia* status of the flies, suggesting they reflect sexual dimorphism in antibacterial defence and not to sex-specific effects of *Wolbachia* (Sex × Wolbachia interaction Fi,i86=0.10, p=0.758).

**Figure 2.**
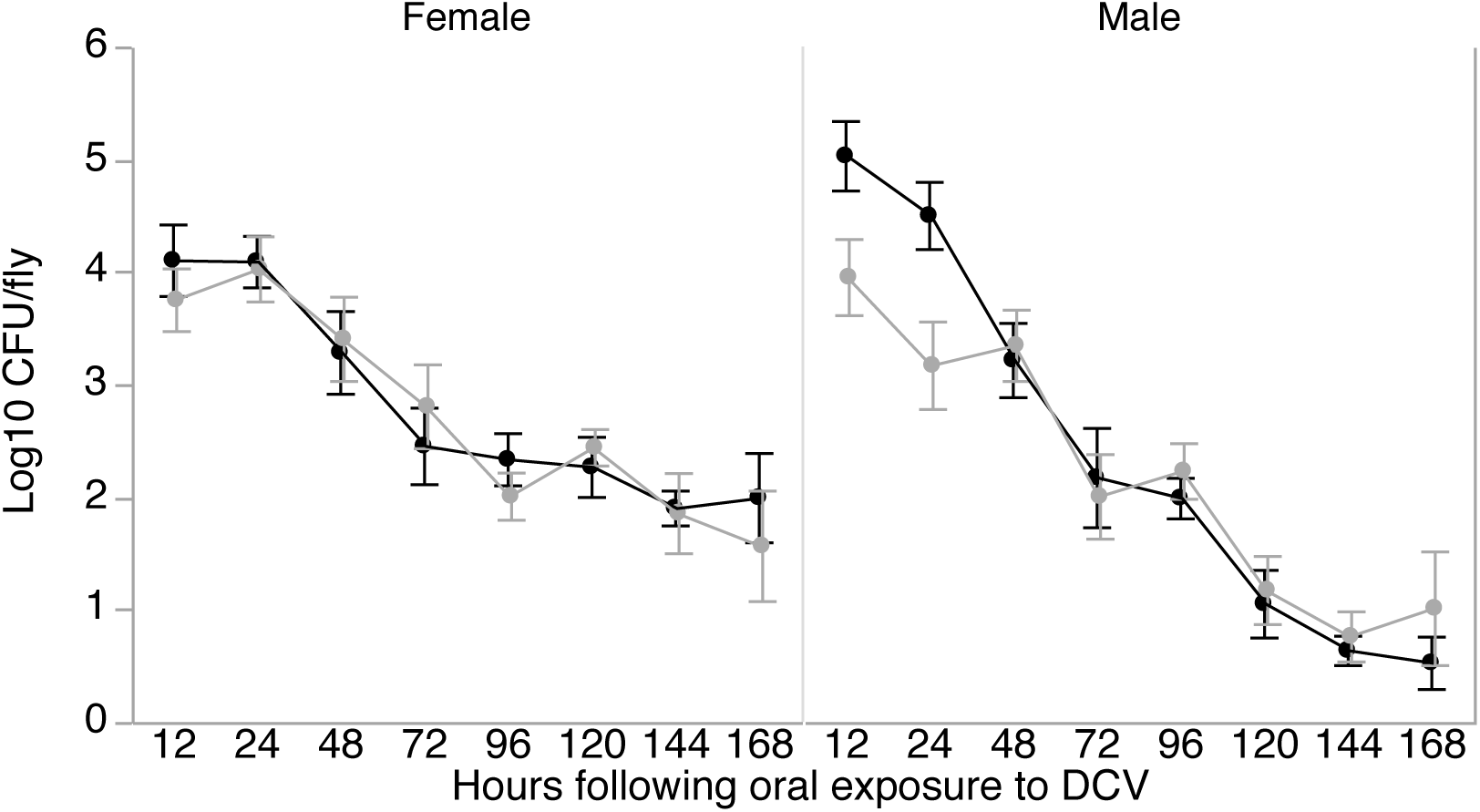
Within-host microbe loads. The number of viable within-host CFUs was quantified in 5-7 individual live flies following 12 hours of oral exposure, and then every 24 hours for a week. Males and females are plotted separately for OreR^Wol-^ (black triangles) and OreR^Wol+^ (grey circles) flies. Data shown are means ± SEM.

### Male flies with *Wolbachia* have lower bacterial loads in the early stages of enteric infection

The initial stages of exposure to pathogens can be crucial in determining whether the host controls infection or if a pathogen grows to a point where hosts are killed. Even though we detected no difference in the rate of clearance according to *Wolbachia* status throughout the infection, at 12 and 24-hours post-infection male flies harbouring *Wolbachia* showed significantly lower bacterial CFUs compared to those without *Wolbachia* (Fig 2; Wol+: 3.86±0.22 Log_10_ CFU; Wol-: 4.56±0.22 Log_10_CFU; F_1,20_ = 5.27, p= 0.033). One explanation for the difference in initial microbe loads in males is that *Wolbachia* could cause behavioural changes, such as reduced feeding rate, that result in reduced infection. However, we did not find evidence that the lower CFU numbers seen in *Wolbachia-positive* male flies resulted from lower feeding rates (Fig. S1). Another potential explanation for the difference in initial microbe loads in males is a *Wolbachia*-mediated antimicrobial response. Whole fly homogenate added to growing cultures of PA14 (in both liquid and solid growth medium) showed greater antimicrobial activity in homogenates of *Wolbachia* positive flies (Figure S2 and S3).

### *Wolbachia-positive* flies show increased expression of IMD pathway genes during the early stages of enteric infection

Given the preliminary evidence of increased antimicrobial activity in *Wolbachia*-positive males (Figure S2 and S3), we decided to test for differences in the expression of antimicrobial immune pathways. While previous work has found no effect of *Wolbachia* on the expression of immune genes (Wong et al., 2011), or on bacterial clearance(Rottschaefer and Lazzaro, 2012), this has not been tested in the context of enteric bacterial infection. Other studies in *Wolbachia*-free flies have demonstrated that the IMD pathway plays an active role in the response to enteric bacterial infection (Buchon et al., 2014, 2009; Kuraishi et al., 2011). We therefore tested whether flies carrying *Wolbachia* showed increased expression of genes involved in IMD-mediated antimicrobial immunity.

In *Wolbachia*-positive females, we observed a significant increase in expression (relative to uninfected females) of two IMD pathway receptor genes-*pgrp-lc* (p = 0.0002) and *pgrp-le* (p = 0.004) at 96-hours post-infection (Fig. 3). In PA14-infected males, however, carrying *Wolbachia* resulted in a slight decrease in *pgrp-lc* expression relative to uninfected males (p = 0.03), although this difference was transient and only observed at 24-hours post-infection (Fig. 3). Overall there appears to be little effect of *Wolbachia* on the expression of either receptor gene in male flies (Fig 3). We observed a significant 3 to 4-fold increase in the expression of the antimicrobial peptide (AMP) gene *attacinA* (*attA*) in *Wolbachia*-positive males at both 24 hours (p = 0.002) and 96 hours (p < 0.001) post-infection. We did not detect any difference in the expression of this AMP gene in male flies that were free of *Wolbachia*, or in female flies, regardless of their *Wolbachia* status (Fig. 3). These results suggest that the initial difference in clearance between Wol+ and Wol-male flies could at least in part be due to *Wolbachia*-mediated up-regulation of the AMP *attacinA*.

**Figure 3.**
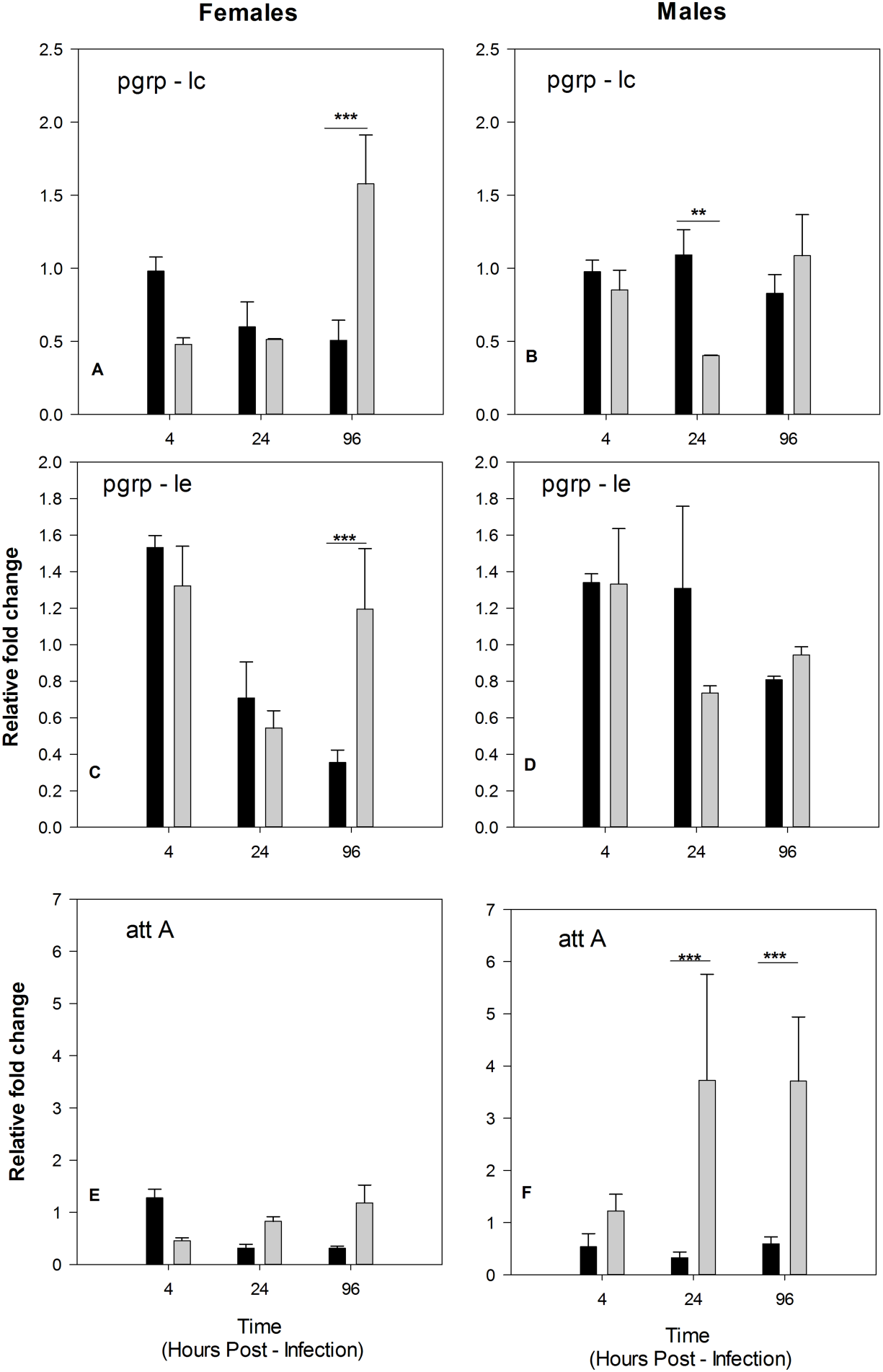
Gene expression of IMD pathway genes. Figures show gene expression relative to *rp49*control gene in infected flies relative to uninfected flies. Mean of 3 biological replicates ± SE.

### *Wolbachia* contributes to increased disease tolerance in male flies

The data we describe above reveals interesting differences in the way male and female flies fight enteric bacterial infection. Male flies are able to clear infection almost completely, while female flies stop clearing infection after 96 hours and maintain a stable bacterial load following that same time period, which suggests that male flies are better than females at clearing enteric PA14 infection **(Fig. 2)**. We could therefore expect females to pay a survival cost due to higher bacterial loads but instead we find that female flies have a similar survival probability to males, especially for flies that are *Wolbachia* negative **(Fig. 1)**. This suggests that females are better able than males to tolerate *P. aeruginosa* enteric infection because they are able to maintain a similar level of health to females, while tolerating higher bacterial loads (Ayres and Schneider, 2012; Medzhitov et al., 2012; Râberg et al., 2009). Male flies, however, showed a marked increase in survival when they were *Wolbachia* positive compared to *Wolbachia* negative males **(Fig. 1)**, even though the rate at which both groups clear infection appear identical **(Fig. 2)**. This suggests increased tolerance in males mediated by the presence of *Wolbachia.* In females, the survival benefit of *Wolbachia* appeared to be minimal, suggesting that *Wolbachia*-mediated tolerance could be sex-specific.

To better assess these differences in disease tolerance mediated by sex and *Wolbachia* status, we plotted the relationship between host health and microbe load for matching time-points (Fig. 4). In all cases, these data were better described by a nonlinear 4-parameter logistic model than a linear model (Table S1). The 4-paraemter logistic model is useful to compare how its maximum (reflecting health in the initial stages of infection), baseline (reflecting the lowest survival reached during the experiment), inflection point (the point at which fly survival reached halfway between the baseline and maximum), and the growth rate (reflecting the rate at which fly survival plummets) vary according to host sex and *Wolbachia* status. Each of these parameters may reflect distinct mechanisms of damage repair involved in host infection tolerance, so they are useful for further exploration of tolerance mechanisms (Ayres and Schneider, 2012; Vale et al., 2016). In female flies, the logistic model explained about a quarter of the variance (R^2^=0.24), and a formal parallelism test found that the curves did not show significantly different shapes (F_3,72_=0.886, p=0.452). In male flies, the 4-parameter logistic model explained over half the variance (R^2^ =0.57), and a formal parallelism test revealed significant differences n the shapes of these two tolerance curves between *Wolbachia*-positive and *Wolbachia*-negative males (F_3,72_ = 2.98, p=0.037). These differences arise not only to the consistently lower maximum and baseline survival in Wolbachia-negative males regardless of microbe load (Figure 4), but also due to differences in the inflection point of each curve which occurs later in the infectious period in Wolbachia-positive male flies (Figure 4).

**Figure 4.**
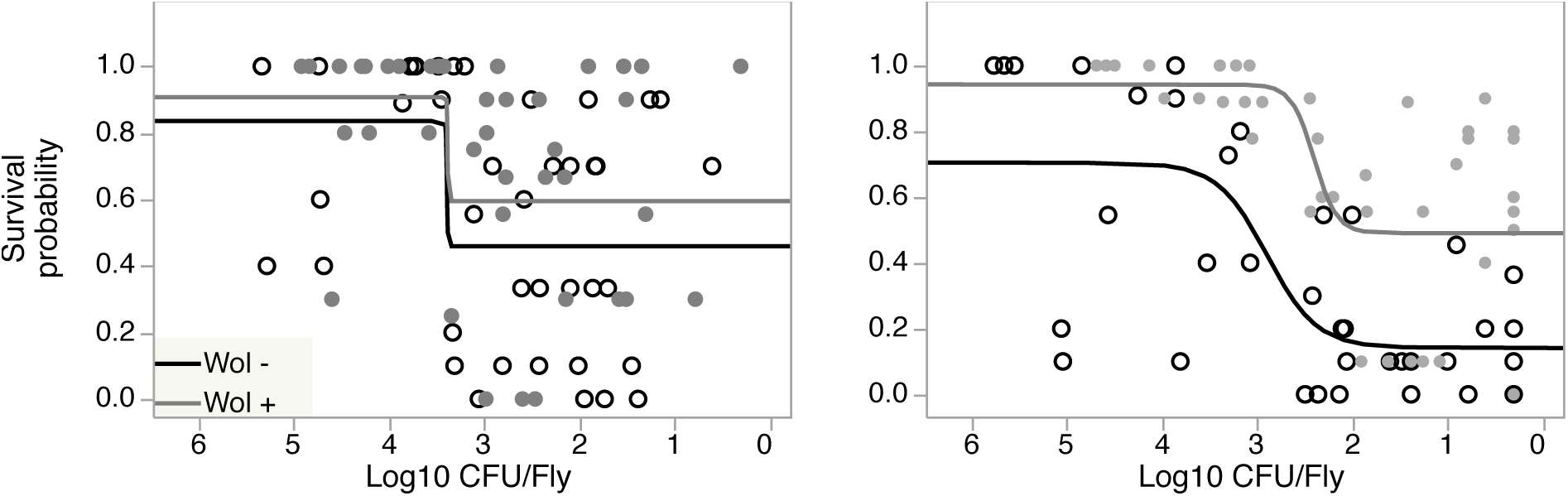
Disease tolerance. To measure tolerance we analysed the relationship between host health and microbe loads. For each time point, we plot the survival probability (as a measure of health) against the microbe load (number of CFU per fly) for 5 biological replicates per sex and *Wolbachia*combination. Here we show the fit of a 4-parameter Logisitic model to the data (see Table S1 for model fits). The X axis is reversed to read from beginning to the end of the infection (only clearance occurred.)

### *Wolbachia* is associated with higher expression of a ROS detoxification gene in males and epithelial repair genes in females during enteric infection

Damage limitation mechanisms such as those involved in the response to oxidative stress, epithelial renewal and damage repair improve host health during infection while not acting directly to eliminate pathogens. They are therefore putative mechanisms of disease tolerance (Ayres and Schneider, 2012; Vale et al., 2014) and have previously been shown to be up-regulated during enteric bacterial infection (Buchon et al., 2009). Given the differences we observed in the ability to tolerate enteric bacterial infection (Fig. 4) we hypothesized that male and female flies may differ in the expression of such genes according to their *Wolbachia* status.

In male flies, enteric infection with PA14, led to increased expression of *gstD8*-involved in ROS detoxification (Buchon et al., 2009; Ha et al., 2005) – which was significantly higher at 96-hours post-infection in those harbouring *Wolbachia*, while no difference flies was observed in female *gstD8* expression according to *Wolbachia* status.

Since oral infection results in damage to insect guts (Buchon et al., 2014), we also measured the expression of two genes involved in epithelial renewal and damage repair(*gadd45* and CG32302) (Buchon et al., 2009). Both genes showed a significant increase in expression in *Wolbachia*-positive females. *Gadd45* expression was marginally higher in *Wolbachia*-positive females compared to those with out *Wolbachia* at 24-hours postinfection (p = 0.045), but this differnce increased by 96-hours post-infection (p < 0.001). CG32302 expression was only transiently differentially expressed in *Wolbachia*-positive females at 24-hours post-infection. Wolbachia-negative males showed a significantly higher expression relative to *Wolbachia*-positive males of both *gadd45* (p = 0.014) and CG32302 (p = 0.004) at 24 hours post-infection, although this difference was no longer observed by 96-hours post-infection.

## Discussion

During the last decade, it has become well established that endosymbionts like *Wolbachia* play a key role in conferring protection from pathogens in their insect hosts (Brownlie and Johnson, 2009; Hamilton and Perlman, 2013; Karyn N. Johnson, 2015). In its natural host *Drosophila, Wolbachia*-mediated protection is especially evident during viral infections (Chrostek et al., 2013; Hedges et al., 2008; Martinez et al., 2014; Teixeira et al., 2008), but protection from bacterial pathogens in *Drosophila* had not been demonstrated to date (Rottschaefer and Lazzaro, 2012; Wong et al., 2011). Here we show that the route of infection is important for *Wolbachia*-mediated protection from bacterial infection. We find that *Wolbachia* can protect *Drosophila* from enteric bacterial infection by eliciting a combination of resistance and disease tolerance mechanisms, and that these protective effects are sexually dimorphic.

### The route of infection matters

The role of *Wolbachia* in protecting hosts from infection, either by increasing resistance or tolerance, is known in *Drosophila*-virus interactions, but previous work testing for antibacterial protection in *Drosophila* did not find a significant effect of *Wolbachia* (Rottschaefer and Lazzaro, 2012; Wong et al., 2011). Typically flies in previous studies were inoculated by intra-thoracic pricking or injection, and therefore experienced a systemic infection. In the wild however, infections are more likely to be acquired through the fecal-oral route (during feeding on decomposing fruit), with most pathogens colonising the gut before being shed through the faeces. *Drosophila-Wolbachia* interactions would therefore have co-evolved mainly under selection by pathogen infection in the gut, and any antibacterial protection that may have evolved as a consequence would not be expected to manifest during a highly virulent systemic infection (Liehl et al., 2006; Martins et al., 2013). Further, if *Wolbachia*-mediated protection is especially efficient in the fly gut, the damage caused by a generalised systemic infection could overwhelm any localised protection by *Wolbachia*, which could explain the lack of observed protection in previous studies of systemic bacterial infection in *Drosophila*. Future studies of host resistance and tolerance should therefore favour natural routes of infection in order to gain a more realistic picture of the mechanisms that hosts have evolved to fight infection.

### *Wolbachia*-mediated protection is a combination of pathogen clearance and damage limitation

The mechanisms underlying *Wolbachia*-mediated protection are presently unclear, especially given that the extent of the protection, and whether it acts to increase resistance or tolerance appear to be pathogen specific (Chrostek et al., 2013; Martinez et al., 2014; Teixeira et al., 2008). In mosquitos *Wolbachia* protection appears to be involved in a combination of general immune priming (Rancès et al., 2012), resource competition between *Wolbachia* and infectious agents (Caragata et al., 2013), and the regulation of host genes involved in blocking pathogen replication (Zhang et al., 2013). In the current experiment it is notable that bacterial numbers did not increase throughout the course of the infection, but were cleared at a near exponential rate (Fig. 2). Despite this, flies still died from infection, although *Wolbachia* reduced the mortality rate. One possibility is that most of the damage experienced by the host happens at the early stages of infection, and the fact that *Wolbachia*-positive flies show lower bacterial titres immediately after the exposure period (Fig. 2) may be the reason they also show higher survival later during the infection. This may explain why the male *Wolbachia* positive flies showed a slower rate of mortality, because *attacinA*-mediated clearance of PA14 within the first 24 hours post infection (Fig. 3) would have minimized gut damage caused by pathogen growth.

An alternative, although not mutually exclusive, possibility is that most of the damage that causes fly death arises from immunopathology, as an indirect side effect of mechanisms that clear pathogens. For example, a common and broad response to infection in insects is the activation of pathways that result in the production of reactive oxygen species (ROS)(Buchon et al., 2014; Ha et al., 2005; Zug and Hammerstein, 2015). ROS production is tightly regulated (Buchon et al., 2013; Ha et al., 2005), and only activated in response to pathogenic and not commensal bacteria (Lee et al., 2013). Previous work has shown that both mosquitoes (Pan et al., 2012) and flies (Wong et al.,2015) harbouring *Wolbachia* show higher ROS levels, but also that in some cases ROS production can lead to oxidative stress, DNA damage (Brennan et al., 2012) and damage to the fly gut epithelium (Buchon et al., 2014, 2013). We therefore hypothesised that mechanisms involved in detoxifying ROS during enteric infection with PA14 may underlie the differences in survival between flies with and without *Wolbachia* (Figure 1). We chose to measure the expression of *gstD8*, involved in ROS detoxification, because it was previously shown to be up-regulated during enteric infection in *Drosophila* with another bacterial pathogen, *Erwinia carotovora* (Buchon et al., 2009).

We found that the expression of *gstD8* was elevated in *Wolbachia*-positive males, but not female flies following 96-hours of oral exposure to *P. aeruginosa.* This pattern of expression is consistent with the increased survival observed in *Wolbachia*-positive males compared to males without the endosymbiont (Figure 1b).

In addition to this detoxification response, we also measured the expression of genes involved in tissue damage repair(*gadd45*) and a component of the peritrophic matrix (*CG32302*), a protective barrier in the fly gut (Lehane, 1997). In males, the presence of *Wolbachia* did not result in an increase in these genes within 96 hours of oral exposure to PA14, but females carrying the endosymbiont showed significantly higher expression than *Wolbachia*-negative flies of *gadd45.* This shows that *Wolbachia* induces different damage limitation mechanisms in males (ROS detoxification) and females (tissue damage repair). We also observed transient increases in the expression of CG32302, another component of gut renewal, in *Wolbachia*-positive females at 24-hours post-infection). There was also a transient increase in expression at 24-hours post-infection of *gadd45* and *CG32302* in *Wolbachia*-negative males (Figure 5). We interpret these increases as a response to increased damage to gut tissue cause by the 10-fold higher bacterial loads in these flies after 24 hours (Fig. 2), which was avoided in *Wolbachia*-positive males by *attacinA*-mediated clearance.

**Figure 5.**
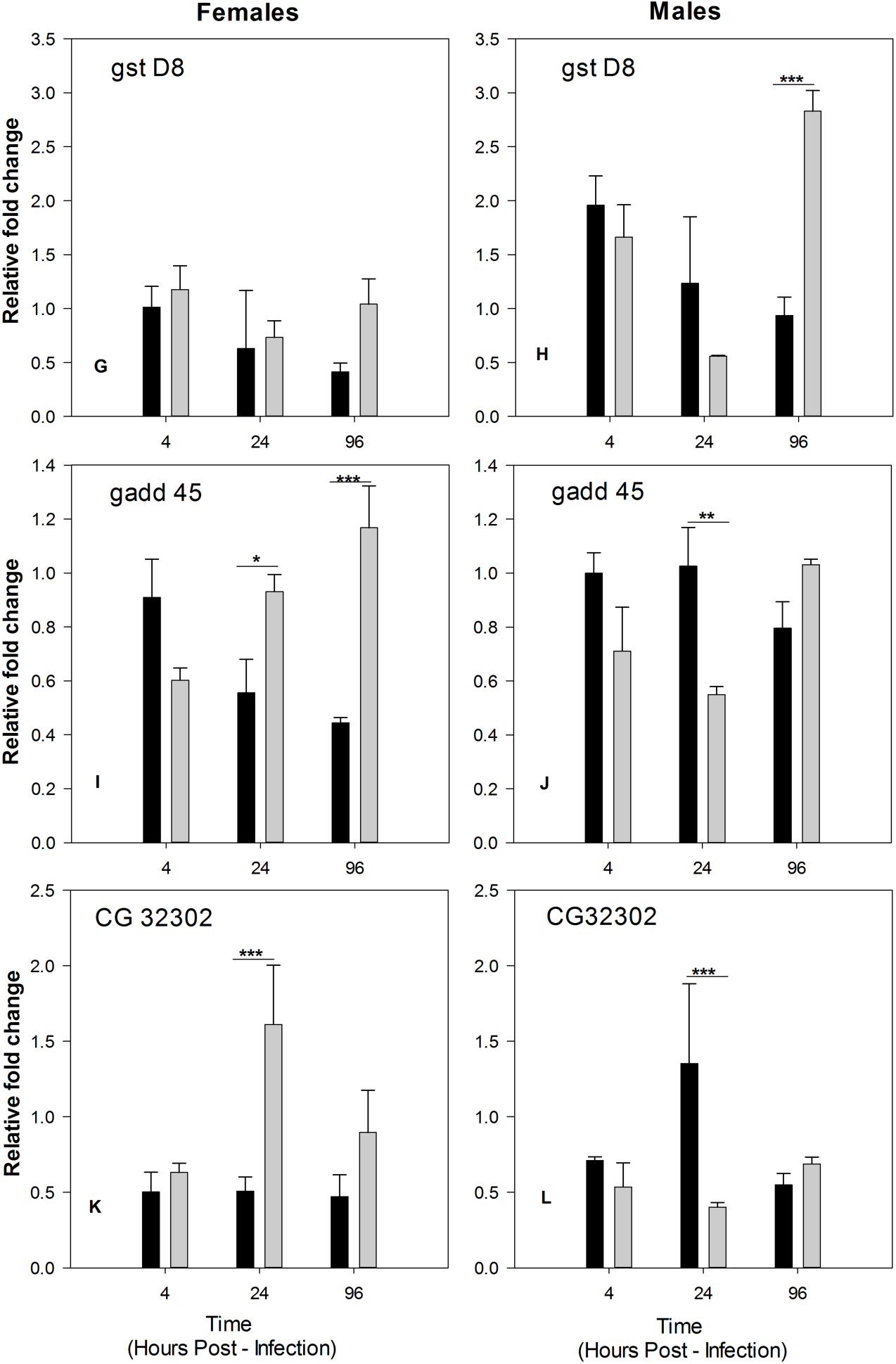
Gene expression of IMD pathway genes. Figures show gene expression relative to *rp49*control gene in infected flies relative to uninfected flies. Mean of 3 biological replicates ± SE.

While previous work found no difference in genome-wide expression levels in adult *Drosophila* with or without *Wolbachia* (Teixeira, 2012), and only mild up-regulation of immune genes has been reported in *Drosophila* cell lines that are transiently infected (Xi et al., 2008), our gene expression results indicate that reducing immunopathology underlies *Wolbachia*-mediated protection from enteric bacterial infection. We are only beginning to understand the complex sequence of events that occur during gut infection in *Drosophila* (Lemaitre and Miguel-Aliaga, 2013), which not only consist of antimicrobial defence, but a multifaceted response that includes stress response, DNA damage repair, the renewal of damaged epithelial cells and gut structure, and the maintenance of efficient metabolism (Buchon et al., 2010, 2009; Kuraishi et al., 2011).

### Sexual dimorphism in resistance and tolerance

The differences in gene expression we describe above reflect two major forms of defence against infection: mechanisms that eliminate pathogens to reduce infection loads, leading to resistance, and mechanisms that limit the damage caused by infection without directly targeting the number of pathogens, leading to disease tolerance (Ayres and Schneider, 2012; Medzhitov et al., 2012; Råberg et al., 2009; Vale et al., 2014). While the majority of work on bacterial and viral infections in *Drosophila* (and most other hosts) has historically focused on mechanisms that eliminate and clear pathogens (Buchon et al., 2014; Obbard et al., 2009; Zambon et al., 2006), the role of mechanisms that limit damage to increase host tolerance is increasingly recognised (Ayres and Schneider, 2012; Medzhitov et al., 2012; Råberg et al., 2009; Soares et al., 2014; Vale et al., 2014). For example recent work has highlighted how tissue damage repair (Jamieson et al., 2013; Soares et al., 2014), immune regulation (Merkling et al., 2015; Sears et al., 2011), and detoxification (Gozzelino et al., 2012; Pamplona et al., 2007) all play a role in enhancing disease tolerance. Variation in disease tolerance is common (Adelman et al., 2013; Howick and Lazzaro, 2014; Råberg et al., 2007; Vale and Little,2012), and may arise from genetic differences in the physiological mechanisms that promote greater tolerance (Råberg et al., 2007), variation in host nutritional states (Howick and Lazzaro, 2014; Sternberg et al., 2012; Vale et al., 2011), and host gut microbiota (Yilmaz et al., 2014). Here we also find evidence for variation in disease tolerance associated with the presence of *Wolbachia*, and we find that these effects vary according to host sex.

*Wolbachia*-mediated protection against viral infections, such as Drosophila C Virus (DCV) acts by reducing viral replication (or increasing the host’s ability to clear infection), suggesting that *Wolbachia* increases resistance against DCV (Hedges et al., 2008; Martinez et al., 2014; Teixeira et al., 2008). In other viral infections, for example Flock House Virus (FHV), flies harbouring *Wolbachia* appear to become more tolerant to infection, showing increased survival without any change in viral titres (Chrostek et al., 2013; Teixeira et al., 2008). Our results show that *Wolbachia* affects both resistance and tolerance to *P. aeruginosa* enteric infection and we found interesting differences between sexes in these responses. Without *Wolbachia* females were more tolerant than males, while males flies became more tolerant when carrying *Wolbachia*. While males and females are generally susceptible to the same pathogens, sexual dimorphism in the immune response is apparent in a wide range of species (Duneau and Ebert, 2012; Marriott and Huet-Hudson, 2006; Zuk and McKean, 1996), and is documented for all classes of viral, bacterial, fungal and parasitic infections [see (Cousineau and Alizon,2014) for a review). In invertebrate hosts, and especially in *Drosophila*, most studies investigating the ability to resist or tolerate bacterial and viral infections have focused primarily on the underlying immune mechanisms (Ayres and Schneider, 2012; Buchon et al., 2014; Kemp and Imler, 2009; Neyen et al., 2014; Schneider et al., 2007), and typically these studies have not focused on sexual differences in these mechanisms [but see (Vincent and Sharp, 2014)). Our results, together with a large body of work on immune sexual dimorphism (Nunn et al., 2009), show that resistance and tolerance mechanisms are likely to vary between males and females.

### Evolutionary and epidemiological implications of sex-specific resistance and tolerance

Variation in resistance and tolerance will directly impact the pathogen loads within hosts (Gopinath et al., 2014; Lass et al., 2013; Susi et al., 2015; Vale et al., 2013), and as a result, sexual dimorphism in these responses could generate potentially important heterogeneity in pathogen spread and evolution (Cousineau and Alizon, 2014; Duneau and Ebert, 2012; Gopinath et al., 2014; Vale et al., 2014). Given that *Wolbachia* are estimated to be highly prevalent in insect populations (Weinert et al., 2015), it is intriguing to consider the potential effects of sexual dimorphism in resistance and tolerance in populations. Theoretical work shows that markedly different evolutionary outcomes for the pathogen are expected when sexual dimorphism in resistance and tolerance is present (Cousineau and Alizon, 2014). One reason is that the mortality rate of males and females will vary due to dimorphism in resistance and tolerance, which in itself will affect the evolutionary trajectories of pathogens (because it will bias infection towards the one sex more than another). Further experimental work is currently needed to test how pathogens are likely to evolve under different host sex ratios, especially when sexual dimorphism in resistance and tolerance is present. Our work suggests that bacterial oral infection in flies benefiting from sex-specific *Wolbachia*-mediated tolerance would offer a useful model system to address these questions.

## Materials And Methods

### Fly stocks

Experiments were carried out using long-term lab stocks of *Drosophila melanogaster* Oregon R (OreR). This line was originally infected with *Wolbachia* strain wMel, (OreR^Wol+^). To obtain a *Wolbachia*-free line of the same genetic background (OreR^Wo-^), OreR^Wol+^ flies were cured of *Wolbachia* by rearing them on cornmeal Lewis medium supplemented with 0.05 mg/ml tetracycline. This treatment was carried out at least 3 years before these experiments were conducted, and the *Wolbachia* status of both fly lines was verified using PCR with primers specific to *Wolbachia* surface protein (wsp): forward (5’-3’): GTCCAATAGCTGATGAAGAAAC; reverse (5’-3’):CTGCACCAATAGCGCTATAAA. Both lines were kept as long-term lab stocks on a standard diet of cornmeal Lewis medium, at a constant temperature of 18±1ºC with a 12-hour light/dark cycle. Flies were acclimatised at 25ºC for at least two generations prior to experimental infections.

### Bacterial cultures

Infections were carried out using the *P. aeruginosa* reference strain PA14, which has been shown to have a very broad host range (He et al., 2004; Mikkelsen et al., 2011). To obtain isogenic PA14 cultures, a frozen stock culture was streaked onto fresh LB agar plates and single colonies were inoculated into 50 mL LB broth and incubated overnight at 37 ºC with shaking at 150 rpm. Overnight cultures were diluted 1:100 into 500 mL fresh LB broth and incubated again at 37 ºC with shaking at 150 rpm. At the mid log phase (OD_600_ = 1.0) we harvested the bacterial cells by centrifugation at 8000 rpm for 10 min, washed the cells twice with 1xPBS and re-suspended the bacterial pellet in 5% sucrose. The final inoculum was adjusted to OD_600_ = 25, and this was the bacterial inoculum used for all flies inoculated orally (enteric infection).

### Enteric and systemic *P. aeruginosa* infection

For systemic infection, flies were pricked at the pleural suture with a needle dipped in a mid log phase (OD_600_ = 1.0) PA14 culture, grown as described above. Control flies were pricked with a needle dipped in sterile LB broth. For the oral exposure (enteric infection), the concentrated PA14 inoculum (OD_600_ = 25) was spotted onto a sterile filter paper (80 μ1/ filter paper), and placed onto a drop of solidified 5% sugar Agar inside the lid of a 7ml Bijou tube. For the uninfected control treatment, filters received the equivalent volume of 5% sucrose solution only. All filter papers were allowed to dry for 20 to 30 minutes at room temperature. We prepared one of these “inoculation lids” for each individual fly. Two to four day-old flies were sex sorted and transferred individually to empty plastic vials: 180 (90 male and 90 female) OreR^Wol+^, and 180 (90 male and 90 female) OreR^Wol-^. Following 2-4 hours of starvation, flies were transferred individually to 7 ml Bijou tubes, and covered with previously prepared “inoculation lids” containing a filter paper soaked in PA14 culture. Flies were left to feed on the bacterial culture for approximately 12 hours at 25 ºC. Following this period, we sacrificed 6 exposed and 2 control flies and counted CFUs by plating the fly homogenate in *Pseudomonas* isolating media (PAIM). The remaining flies were transferred to vials containing 5% sugar agar and incubated at 25 ºC.

### Quantification of within-host bacterial loads

Following the initial 12-hour exposure, every 24 hours we randomly sampled 5 to 7 live flies per sex and *Wolbachia* status and quantified the microbe loads present inside the flies. Briefly, a single fly was removed from the vial and transferred to 1.5 mL microcentrifuge tubes. To guarantee we were only quantifying CFUs present inside the fly, and not those possibly on its surface, each fly was surface sterilized by adding 75% ethanol for 30-60 seconds (to kill the outer surface bacterial species). Ethanol was discarded and flies were washed twice with distilled water. Plating 100 μL of the 2^nd^ wash in LB agar confirmed this method was efficient in cleaning the surface of the fly (no viable CFUs were detected). Each washed whole fly was placed in 1 mL of 1X PBS in a 1.5-mL screw-top microcentrifuge tube, centrifuged at 5000 rpm for 1 min and the supernatant was discarded. 200 μL of LB broth was then added to each tube and the flies were thoroughly homogenised using a motorised pestle for 1 min. A 100 μL aliquot of homogenate was taken for serial dilution and different dilutions were plated on PAIM agar plates, incubated at 37 ºC for 24 – 48 h and viable CFUs were counted.

### Survival assays

We carried out separate experiments to measure how the presence of *Wolbachia* affected fly mortality during either enteric or systemic infection, with identical fly rearing and bacterial cultural conditions as those described above. For each survival assay (enteric or systemic infection routes), two-to-four day-old flies were sexed and exposed in groups of 10 flies to PA14, as described above. For each combination of male or female OreR^Wol+^ and OreR^Wol-^ line, we set up fifty flies, with 10 flies per vial. The flies that died from infection was recorded every approximately every hour until all flies had died (systemic infection), or every 24 hours for up to 8 days (enteric infection).

### Statistical analyses of host survival and microbe loads

Fly survival was analysed using a Cox proportional hazards model to compare survival rates, with fly ‘Sex’, ‘Infection status’, and *‘Wolbachia* status’ and their interactions as fixed effects. The significance of the effects was assessed using likelihood ratio tests following a χ^2^distribution. For flies that were exposed orally to DCV, we compared between pairs of treatments (control vs. infected or with and without *Wolbachia*) using Cox risk ratios.

In orally infected flies, changes in the bacterial load within-hosts were analysed with a linear model with Log CFU as the response variable, and fly ‘Sex’,*‘Wolbachia* status’, and ‘Time (DPI)’ as fixed effects. Differences in the mean bacterial load of flies immediately following oral exposure (at 12 hours post-infection; Fig S1) were also analysed separately using a linear model with Log CFU as the response variable and ‘Sex’ and *‘Wolbachia* status’ as fixed effects. In all models all effects and their interactions were tested in a fully factorial model, and models were simplified by removing non-significant interaction terms. All analyses were conducted in JMP 12 (SAS).

### Analysis of disease tolerance

Disease tolerance is defined as the ability to maintain health relative to changes in microbe loads during an infection(Ayres and Schneider, 2012; Medzhitov et al., 2012; Råberg et al., 2009). It is possible to analyse tolerance as the time-ordered health trajectory of a host as microbe loads change (Doeschl-Wilson et al., 2012; Lough et al., 2015; Schneider, 2011). To assess sex‐ and *Wolbachia*-mediated differences in how sick a fly gets for a given pathogen load (tolerance) during the course of the infection, for each time point we took the survival probability (as a measure host health) and PA14 CFUs present within the flies (as a measure of microbe load) for 5 replicate flies in each sex/*Wolbachia* combination. While tolerance is commonly described as a linear reaction norm (Lefèvre et al., 2011; Råberg et al., 2007; Simms, 2000), many other functional forms are possible, including non-linear relationships (Vale et al., 2014). To assess the form of the health/microbe relationship, we fit linear and non-linear 4-parameter logistic model separately to the time-matched survival/microbe load plots. In all cases, the 4 parameter logistic model – which is commonly use to asses dose-response curves (Gottschalk and Dunn, 2005) – outperformed the linear fit (Table S1), and we present only the logistic fit in the results section. To test if these tolerance curves differed with *Wolbachia* status, we tested the parallelism of the models by comparing the error-sums of square for a full model (where each group has different parameters in the logistic model) to a reduced model (where models share all parameters except the inflection point) (Gottschalk and Dunn, 2005).

### Gene expression

We used qRT-PCR to test for differences in the expression of genes known to be involved in either bacterial clearance or in the response to stress and gut damage during enteric infection. Previous work has shown that the IMD pathway and some stress response and damage repair genes are especially important during the fly’s response to enteric bacterial pathogens (Buchon et al., 2009). We therefore investigated the expression of IMD pathway receptor genes(*pgrp – lc* and *pgrp – le*) and the antimicrobial peptide effector gene *attacin A* (attA); *gstD8*, a gene that participates in the detoxification of reactive oxygen species (ROS) produced during microbial immunity in the gut (Ha et al., 2005); *gadd45*, a gene relevant for epithelial renewal, as it is involved in stress response and wound healing in *Drosophila* (Stramer et al., 2008; Takekawa and Saito, 1998); and CG32302, a gene identified as up-regulated during enteric bacterial infection(Buchon et al., 2009). CG32302 has been annotated as a putative component of the peritrophic matrix (Buchon et al., 2009), a protective barrier that separates the gut epithelium from the invading bacteria (Lehane, 1997).

Our aim was to test if the expression of these genes varied in a sex‐ or *Wolbachia-specific* manner in flies that were infected orally. Groups of 5 flies for each sex / *Wolbachia* combination were exposed orally to *P. aeruginosa* infection in triplicate, as described above, and then frozen in TRI reagent at 4, 24 and 96 hours post-infection. Total RNA was extracted from flies homogenised in Tri Reagent (Ambion), reverse-transcribed with M-MLV reverse transcriptase (Promega) and random hexamer primers, and then diluted 1:10 with nuclease free water. The qRT-PCR was performed on an Applied Biosystems StepOnePlus system using Fast SYBR Green Master Mix (Applied Biosystems) with the following PCR cycle: 95°C for 2min followed by 40 cycles of 95°C for 10 sec followed by 60°C for 30 sec. Three biological replicates and two qRT-PCR reactions per replicate (technical replicates) were carried out per sample. Gene-specific primers are reported in Table S1. Changes in gene expression were analysed relative to the expression of *rp49*, an internal control gene. The relative fold-change difference in expression between infected and health control flies was calculated as described in (Livak and Schmittgen, 2001). Briefly:

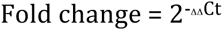

Where, ΔΔCt = [(Ct of Gene A – Ct of Internal control) of Infected sample) – [(Ct of Gene A – Ct of Internal control) of Control sample)

The fold change difference obtained was analysed using 3-way ANOVA with sex (Male, Female), Time (2, 24 and 96 hours) and *Wolbachia* (Wol‐ and Wol+) as fixed factors.

## Acknowledgements

We thank R. Popat for technical assistance, D. Obbard, F. Waldron, T.Little and S. Lewis for helpful discussion and suggestions, and H. Cowan, H. Borthwick for media preparation.

## Funding information

This work was supported by a strategic award from the Wellcome Trust for the Centre for Immunity, Infection and Evolution (http://ciie.bio.ed.ac.uk; grant reference no.

095831), and by a Society in Science – Branco Weiss fellowship (http://www.society-in-science.org), both awarded to P. Vale. P. Vale is also supported by a Chancellor’s fellowship from the School of Biological Sciences

## Author contributions

PFV and RBV conceived the study. RBV, GV, JSJ, KMM and PFV conducted experimental work. GV and PFV analysed the data. PFV wrote the manuscript. SPB and PFV contributed reagents and consumables. All authors commented on the manuscript.

## Supporting Information

***Wolbachia* confers sex-specific resistance and tolerance to enteric but not systemic bacterial infection in *Drosophila***

Radhakrishnan B. Vasanthakrishnan ^1,3§^,^1§^, Gupta Vanika ^1§^, Jonathon Siva-Jothy, Katy M. Monteith^1^, Sam P. Brown^4^, Pedro F. Vale^1,2*^,

**S1 file.** Materials and methods for feeding assay and PA14 growth inhibition assays.

### Feeding assay

All flies used were 0-72 hour old adult Oregon R flies, and Wol‐ and wol+ individuals were raised separately in 6oz bottles on lewis medium at a constant temperature of 25°C on a 12:12 light:dark cycle. Subjects were starved for 2 hours by placing them an empty vial before CO2 anaesthesia was used to separate sexes. While still anaesthetised each fly was placed into an individual 23ml vial containing Blue-dyed Lewis medium (Blue Dye number 1, 0.5g/litre). Between 22-24 individual flies per Sex/Wolbachia combination were set up. All flies were left to feed on the blue dyed medium for 24hours at 25 degrees under 12:12 light:dark cycle. After 24 hours flies were immediately frozen to kill and then decapitated using a scalpel. Decapitation avoids inaccurate absorbance readings due to eye pigments. Each individual fly was then placed in an Eppendorf tube containing 100ul of ice-cold Ringer’s solution, homogenised using a motorised pestle and then centrifuged for 10mins at ~13300g at 20°C. 80 n.l of this supernatant were loaded into a 96-well plate and the blue pigment was measured using a VersaMax microplate reader, recording absorbance at 520nm.

**Fig S1.**
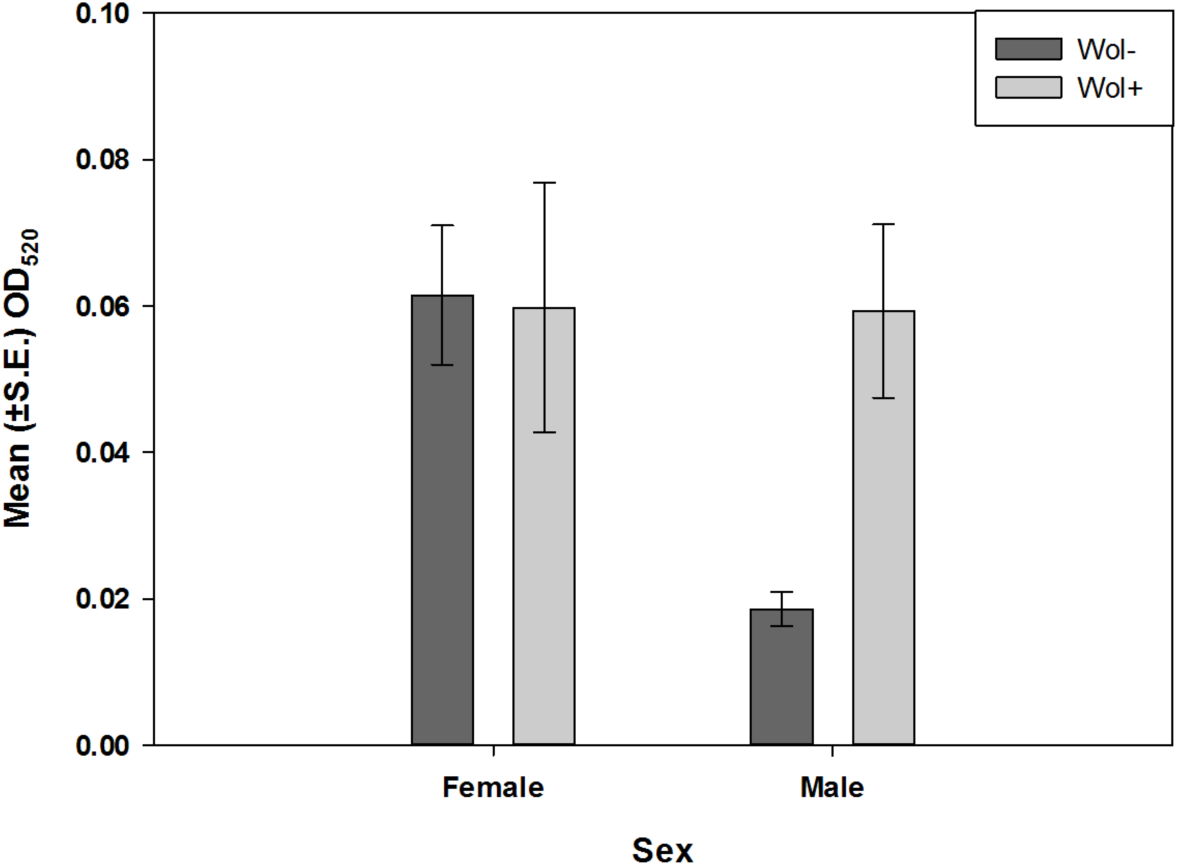
Quantification of food intake. To quantify feeding, individual flies were fed on blue-dyed medium for 24 hours, homogenised and suspended in buffer. The absorbance of this suspension, which is proportional to the amount of food intake (Bashir-Tanoli and Tinsley, 2014), was measured at 520 nm. Data shown are means ± SEM.

### Pseudomonas aeruginosa PA14 inhibition assay

Wol+ and Wol‐ flies were grown in 6 oz plastic bottles on standard Lewis medium under standard laboratory conditions at 25°C, 12:12 Light:Dark cycle. Six days post eclosion, flies were sex separated and divided into cohorts of 10 flies each (n = 10 per sex per line). Flies were anaesthetized using cold anaesthesia. To each sample, 250 μl of LB was added and samples were crushed using an automated pestle and centrifuged at 11,000g for 5 minutes. Supernatant from these samples was used to assay antibacterial response. 100 μl from supernatant was transferred to a well in 96-well plate. PA14 cultures were grown as follows. A single colony was picked from the PA plate and 5 ml of LB medium was inoculated. The culture was grown overnight at 37°C at 200 rpm. 100 μl of the overnight culture was used to seed 5ml of LB. OD was monitored and culture was taken out after it reached OD_600_ = 1. The OD of the culture was adjusted to 0.02. Each well was seeded with 100 μl of this culture and was mixed well with fly homogenate by pipetting. For controls, LB without any bacteria was used. Bacterial growth in 96-well plate was monitored overnight using Thermo Scientific Varioskan Flash. Plate was incubated at 37°C and readings were collected every half an hour intervals for 16 hours. Data obtained was used to obtain lag phase time, growth rate and saturation time for each well using Growth Curve Analysis Tool (GCAT). These data were analysed using ANOVA with fly sex and Wolbachia status as factors.

**Fig S2.**
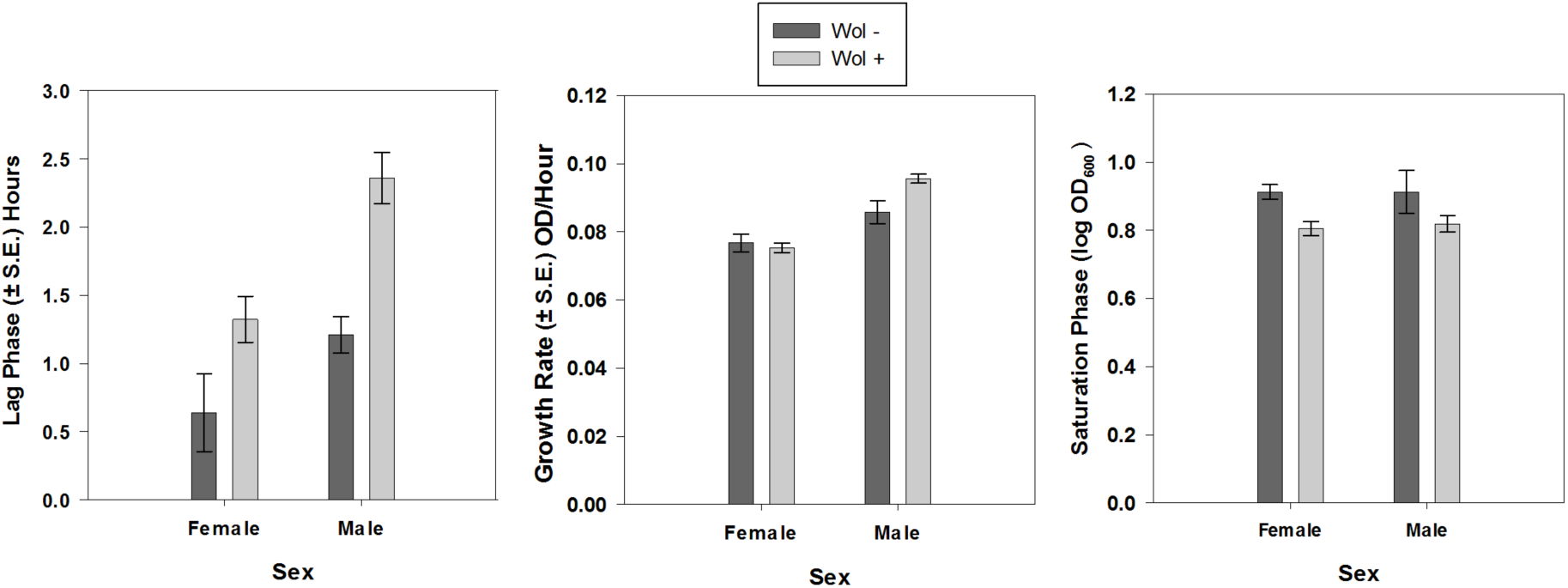
Inhibition of PA14 growth in liquid culture. Single sex groups of *Wolbachia* positive or *Wolbachia* negative flies were homogenised, centrifuged and the supernatant was inoculated into PA14 cultures growing in 96-well plates. Absorbance was measured every 30 minutes.

We also carried out an inhibition zone assay on PA14 grown on a solid medium. Bacterial cultures and fly homogenate were both prepared as described above. When growing the PA14 culture, 100 μl of a 500-fold dilution was used to plate on LB agar plates to obtain a uniform PA14 lawμ After 30 minutes, 100 μl of fly homogenate was put on the plate. Plates were incubated at 37°C for overnight. Clearance zone formed by fly homogenate was measured using Image J and area values were analysed using ANOVA with fly sex and Wolbachia status as factors.

**Fig S3.**
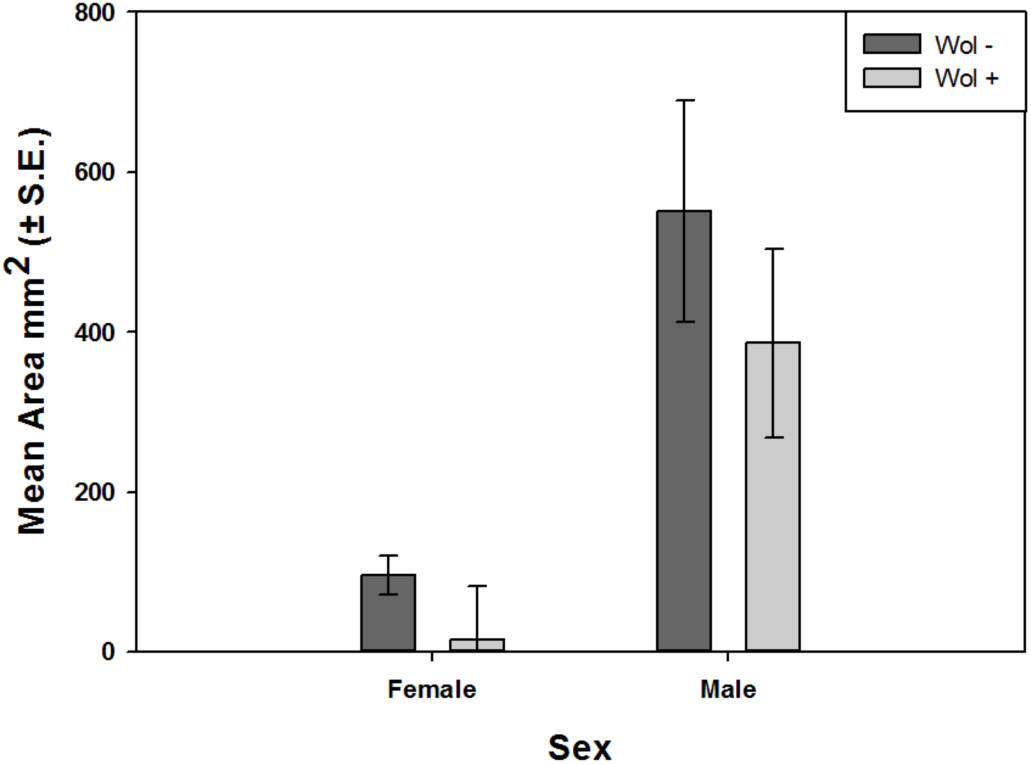
Inhibition zone assay. Single sex groups of *Wolbachia* positive or *Wolbachia* negative flies were homogenised, centrifuged and the supernatant was spotted on a lawn of PA14 overnight. The figure shows the size of the zone where PA14 growth was inhibited. Data shown are means ± SEM.

**Table S1.**
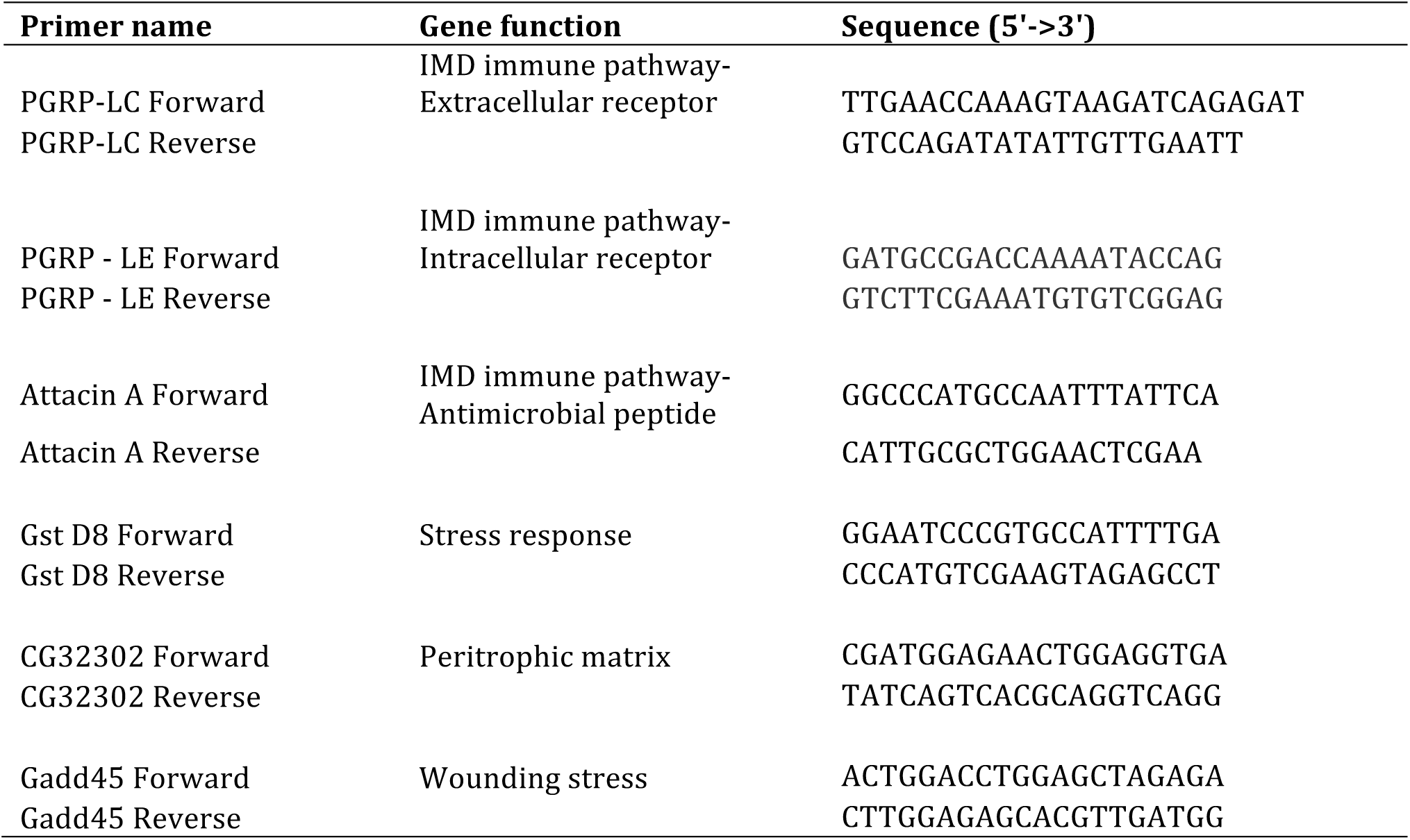
Primer list.

**Table S1.** Fits of non-linear tolerance curves.

| Sex | Model | AICc | AICc Weight | R-Square |
| --- | --- | --- | --- | --- |
| Female | Logistic 4P | 59.83289049 | 0.80614163 | 0.239893273 |
|  | Linear | 62.68315369 | 0.193858371 | 0.110112664 |
| Male | Logistic 4P | 18.97942649 | 0.511289458 | 0.572646097 |
|  | Linear | 19.0697575 | 0.488710542 | 0.516645982 |

